# Polyomavirus T-Antigen Induces *APOBEC3B* Expression using a LXCXE-Dependent and TP53-Independent Mechanism

**DOI:** 10.1101/312595

**Authors:** Gabriel J. Starrett, Artur A. Serebrenik, Pieter A. Roelofs, Brandy Verhalen, Jennifer L. McCann, Matthew C. Jarvis, Teneale A. Stewart, Emily K Law, Annabel Krupp, Mengxi Jiang, John W. M. Martens, Paul N. Span, Ellen Cahir-McFarland, Reuben S. Harris

## Abstract

APOBEC3B is a single-stranded DNA cytosine deaminase with beneficial innate antiviral functions. However, misregulated APOBEC3B can also be detrimental by inflicting APOBEC signature C-to-T and C-to-G mutations in genomic DNA of multiple cancer types. Polyomaviruses and papillomaviruses use dominant oncoproteins to induce APOBEC3B overexpression, perhaps to their own benefit, but little is known about the cellular mechanisms hijacked by these viruses to do so. Here we investigate the molecular mechanism of APOBEC3B upregulation by the polyomavirus large T-antigen. First, truncated T-antigen (truncT) is sufficient for APOBEC3B upregulation and the RB interacting motif (LXCXE), but not the TP53 inhibition domain, is required. Second, upregulated APOBEC3B is strongly nuclear and partially localized to virus replication centers. Third, genetic knockdown of RB1 alone or in combination with RBL1 and/or RBL2 is insufficient to suppress truncT-mediated induction of *APOBEC3B*. Fourth, CDK4/6 inhibition by palbociclib is also insufficient to suppress truncT-mediated induction of *APOBEC3B*. Fifth, global gene expression analyses in a wide range of human cancers show significant associations between expression of *APOBEC3B* and other genes known to be regulated by the RB/E2F axis. These experiments combine to implicate the RB/E2F axis in promoting *APOBEC3B* transcription, yet they also suggest that the polyomavirus RB binding motif has in addition to RB inactivation at least one additional function for triggering *APOBEC3B* upregulation in virus-infected cells.

**IMPORTANCE:** The APOBEC3B DNA cytosine deaminase is overexpresssed in many different cancers and correlated with elevated frequencies of C-to-T and C-to-G mutations in 5’-TC motifs, oncogene activation, acquired drug resistance, and poor clinical outcomes. The mechanisms responsible for APOBEC3B overexpression are not fully understood. Here, we show that the polyomavirus truncated T-antigen (truncT) triggers APOBEC3B overexpression through its RB-interacting motif, LXCXE, which in turn likely enables one or more E2F family transcription factors to promote *APOBEC3B* expression. This work strengthens the mechanistic linkage between active cell cycling, APOBEC3B overexpression, and cancer mutagenesis. Although this mechanism damages cellular genomes, viruses may leverage it to promote evolution, immune escape, and pathogenesis. The cellular portion of the mechanism may also be relevant to non-viral cancers, where genetic mechanisms often activate the RB/E2F axis and APOBEC3B mutagenesis contributes to tumor evolution.

## INTRODUCTION

**G**enetic diversity is key to virus replication, pathogenesis, and transmission, and particularly for escape from adaptive immune responses in vertebrate species (1–3). Each virus has evolved to have an optimized level of genetic diversity for its own unique life cycle, with some viruses having high mutation rates and others much lower mutation rates (4, 5). Multiple processes contribute to the overall constellation of mutations observed in viral genomes, including nucleotide misincorporation by DNA polymerases during replication (4, 2).

Recently, an APOBEC mutation signature has been reported in multiple DNA tumor viruses, including high-risk human papillomavirus (HPV) types and BK polyomaviruses (6–9). Several APOBEC enzymes, including APOBEC3B (A3B), bind 5’-TC dinucleotide motifs in single-stranded DNA and catalyze the hydrolytic conversion cytosine to uracil (10, 11). Left unrepaired, uracil lesions can serve as templates for new DNA synthesis and directly result in C-to-T mutations. Alternatively, if the uracil base is excised by cellular uracil DNA glycosylase 2 (UNG2), then the resulting abasic site becomes non-instructional and may trigger cellular DNA polymerases to insert an adenine opposite the lesion, except REV1 that tends to incorporate either adenine or cytosine. Thus, APOBEC-catalyzed DNA deamination of 5’-TC motifs results in both C-to-T and C-to-G mutations (a signature frequently expanded to include the 3’-nucleobases A and T and referred to in the context of trinucleotide motifs 5’-TCA and 5’-TCT). An additional hallmark of virus mutagenesis by APOBEC enzymes is a bias toward the lagging-strand of DNA replication (12–14). A likely mechanistic relationship with single-stranded DNA replication intermediates is supported by similar correlations in model yeast and *E. coli* experiments (15, 16).

Human cells have the potential to express up to nine active DNA cytosine deaminases (AID, APOBEC1, and A3A/B/C/D/F/G/H) (17–20). Seven of these enzymes prefer 5’-TC motifs in single-stranded DNA, whereas AID uniquely prefers 5’-RC and APOBEC3G (A3G) 5’-CC. A3B is the most likely APOBEC family member to contribute to the mutagenesis and evolution of small DNA tumor viruses because it is specifically upregulated by viral oncoproteins. For high-risk HPV types, the oncoproteins E6 and E7 have been implicated through various pathways (21–24). For polyomaviruses including JC, BK, and Merkel cell (JCPyV, BKPyV, MCPyV), the large T-antigen (TAg) induces A3B upregulation (6). Although these viral proteins bear little similarity at the amino acid sequence level, they have considerable functional overlap including RB inactivation by E7 and TAg and TP53 inactivation by E6 and TAg. Here we investigate the molecular mechanism by which polyomaviruses promote the transcriptional upregulation of *A3B* with results converging on the cellular RB/E2F pathway, which is often deregulated in cancer.

## Results

### Visualization of endogenous APOBEC3B protein in polyomavirus infected cells

A3B induction by polyomaviruses has been shown at the mRNA level by RT-qPCR and at the protein level by immunoblotting in primary renal proximal epithelial cells (RPTECs) (6). To extend these results, immunofluorescent microscopy was used to ask whether polyomavirus infection causes a general pan-nuclear upregulation of A3B enzyme and/or localization to discrete subnuclear regions such as virus replication centers in other relevant cell types. Tenth generation (G10) primary human embryonic kidney (HuK(i)G10) cells were infected with JCPyV (MAD1 strain) and analyzed 7 days later by pulse-labeling 15 minutes with EdU and immunofluorescent microscopy. As shown previously (25), infected cells are distinguished by larger nuclei positive for TAg and EdU (representative images in **Fig. 1A**). Virus replication centers appear as foci that stain even brighter for these two markers. In comparison, A3B is clearly induced in the nuclear compartment of infected cells but not uninfected cells. A3B staining is broadly nuclear and not obviously co-localizing with EdU-positive virus replication foci. TAg and EdU staining have a strong positive correlation signifying active viral DNA replication, and TAg and A3B show a weaker but still significantly positive correlation (**Fig. 1B-C**).

**Figure 1.**
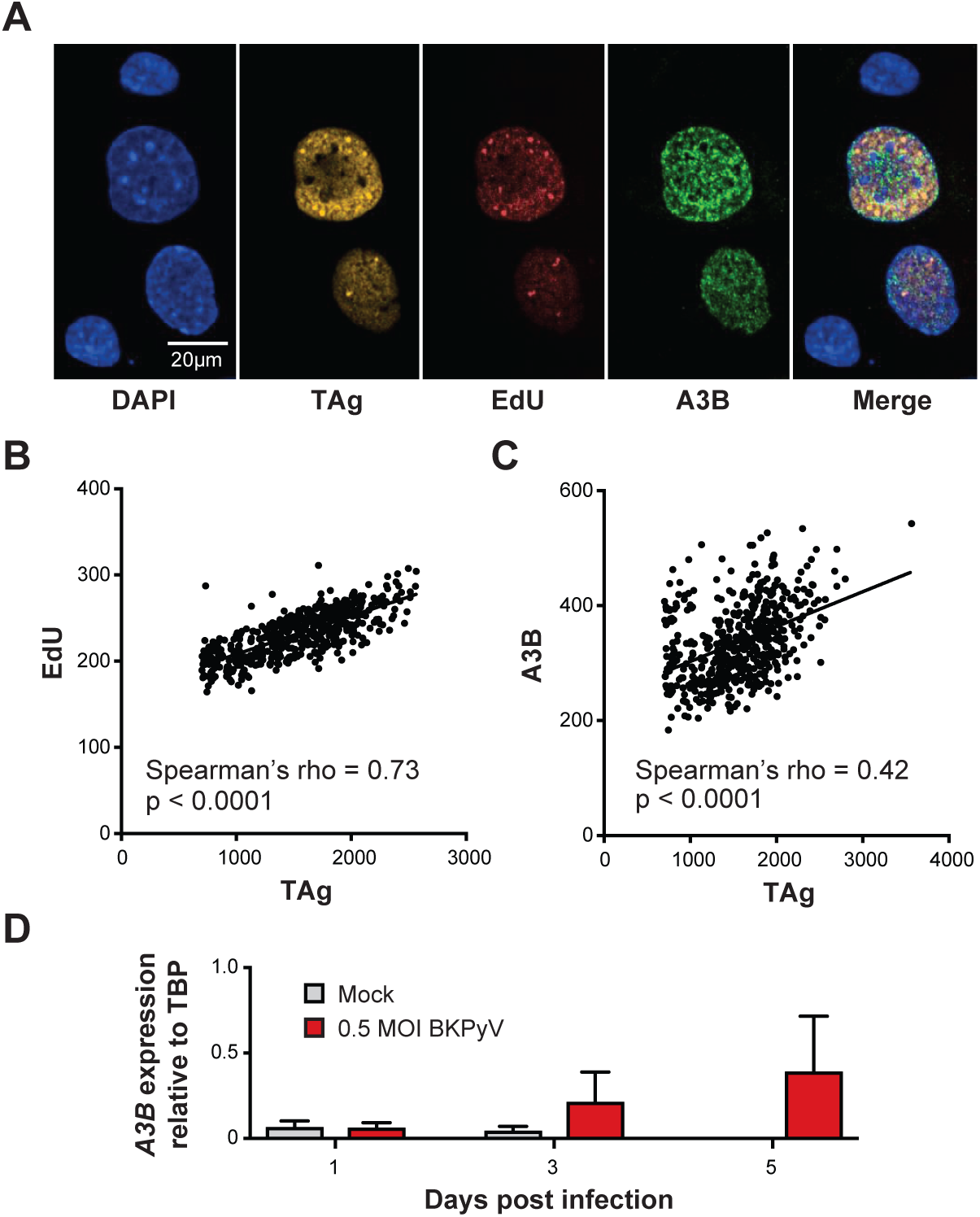
Visualization and quantification of A3B expression in PyV-infected cells. (A) High resolution immunofluorescence of DAPI, A3B, EdU, and TAg in HuK(i)G10 cells infected with MAD1 JCPyV. (B) Quantification and linear correlation of EdU incorporation versus T-antigen intensity from A. (C) Quantification and linear correlation of A3B versus T-antigen intensity from A. (D) Bar graph showing A3B mRNA expression in BK polyomavirus infected (red) and mock infected (gray) MCF10A cells.

To date, many aspects of A3B regulation and function have been determined using normal-like and cancerous mammary epithelial cell lines, because of direct relevance to cancer and a higher capacity for genetic manipulation over primary cells (26, 27). To ask whether TAg induces *A3B* in one of these systems, the normal-like mammary epithelial cell line MCF10A was infected with BKPyV and RT-qPCR was used to quantify *A3B* mRNA levels over time. As above for early passage human embryonic kidney cells, *A3B* expression peaked 3 and 5 days post-infection with mock-treated cells showing no significant changes (**Fig. 1D**). Thus, together with our prior studies (6), polyomavirus appear to possess a conserved and robust capacity for *A3B* induction.

### APOBEC3B upregulation by polyomavirus large T-antigen requires the canonical RB-interacting motif LXCXE

Based on our previous studies (6), the large (L)TAg of multiple different polyomaviruses triggers upregulation of A3B expression in RPTECs. To investigate the LTAg domains responsible for A3B induction, we tested a naturally occurring splice variant of BKPyV LTAg, known as truncT, which lacks the DNA-binding and helicase domains essential for TP53 neutralization (28–30) (schematic in **Fig. 2A**). In parallel, we also assessed derivatives of LTAg and truncT with a disrupted LXCXE motif, which is required for inhibiting the tumor suppressor protein RB, as previously reported using SV40 TAg (31). RPTECs were transduced with lentiviruses expressing an empty multiple cloning site as a negative control, BKPyV LTAg as a positive control, BKPyV truncT, and RB-binding site mutant derivatives, incubated 3 days, and assessed by immunoblotting and fluorescent microscopy. Mock-transduced cells express low levels of A3G, and transduction with empty lentivirus causes a modest increase in this protein and also raises A3B levels to faintly detectable levels (**Fig. 2B**). In contrast, both LTAg and truncT dramatically induce A3B expression, and all induction is eliminated by two amino acid substitutions shown previously to abrogate RB binding in SV40 (LFCHED to LFCHKK) (31) (**Fig. 2B**). Immunofluorescent microscopy images also show truncT-mediated induction of nuclear A3B but not by the RB-binding mutant derivative (**Fig. 2C**). Similar results were obtained for truncT the same RB-binding mutant in the human mammary epithelial cell line MCF10A (**Fig. 2D-E**).

**Figure 2.**
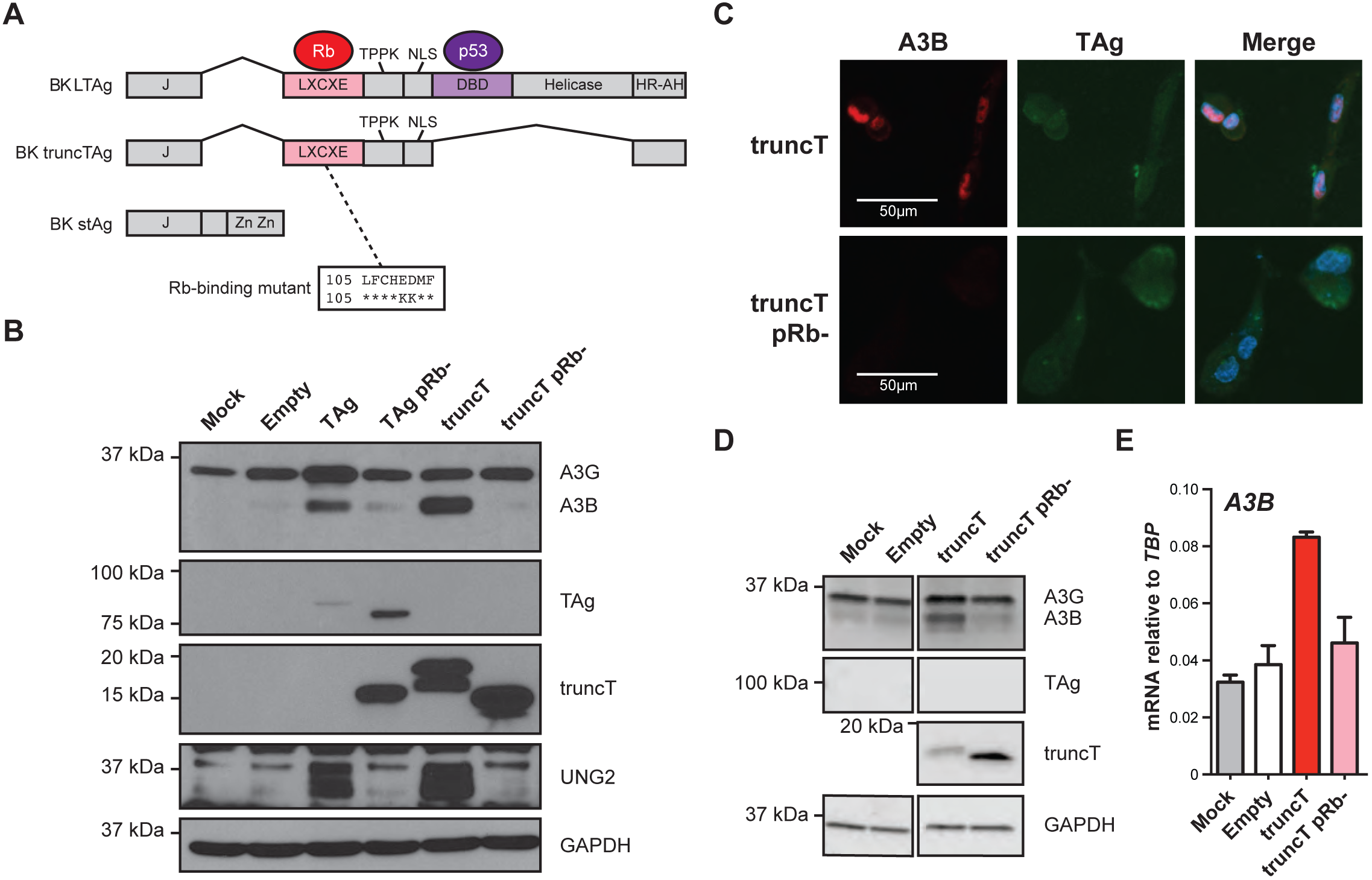
RB-binding domain is necessary for A3B induction by polyomavirus T-antigens. (A) Diagram of BK T-antigen isoforms and LXCXE mutants used in this study. (B) Western blots of RPTE cells transduced with T-antigen mutants and respective A3B expression. (C) Immunofluorescence images for TAg and A3B in T-antigen and mutant transduced RPTECs. (D) Immunoblots of MC10A cells transduced with T-antigen mutants and respective A3B expression. (E) Bar graph showing A3B mRNA expression determined by RT-qPCR for MCF10A cells mock treated (gray) or treated with empty (white), truncT (red) and Rb-binding mutant truncT (pink) lentiviral vectors.

### TP53 inactivation is dispensable for *APOBEC3B* induction

The aforementioned data comparing LTAg and truncT simultaneously implicate RB and demonstrate that TP53 inhibition is not required for A3B induction because truncT completely lacks the TP53 binding domain (**Fig. 2**). To further ask whether TP53 inactivation might influence *A3B* gene expression, we quantified *A3B* mRNA levels in two cell lines that have been used to study A3B regulation, MCF10A and the human estrogen-receptor positive breast cancer cell line MCF-7L [above and (26, 27, 32, 33)]. Each cell line was treated with either DMSO or 5 μM nutlin, which is a drug that protects TP53 from MDM2-mediated degradation (34). As controls, mRNA levels were quantified for three genes repressed by TP53 (*P21*, *MDM2*, and *TP53*) and one gene activated by TP53 (*7LC7A11*). Respectively, the expression of these genes was derepressed or repressed by nutlin treatment (**Fig. 3A-B**). In comparison, neither steady-state nor PMA-induced *A3B* mRNA levels were changed by nutlin (**Fig. 3A-B**). Moreover, Cas9-mediated knockout of *TP53* in MCF10A cells also caused no significant effect on basal or PMA-induced *A3B* expression levels (**Fig. 3C-D**). These data combine to indicate that TP53 has no significant role in the transcriptional regulation of *A3B*, disproving our original hypothesis (21) and conflicting with recently published data (33) (**Discussion**).

**Figure 3.**
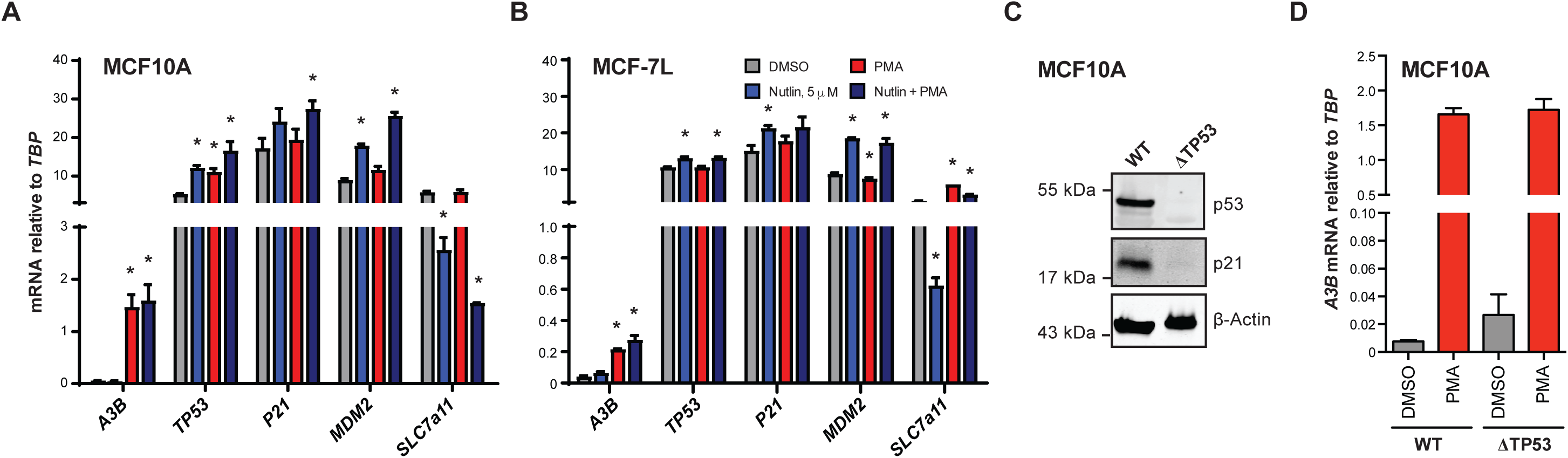
Inactivation of p53 does not affect A3B expression. (A-B) RT-qPCR measurement of relevant genes in MCF10A (A) and MCF7L (B) cells treated with DMSO, 5μM nutlin, PMA, or nutlin + PMA. Statistically significant changes by student’s t-test (p<0.05) are noted by an asterisk. (C) Immunoblot of WT and *TP53* KO MCF10A cell lines. (D) PMA treatment of WT and p53 KO MCF10A cell lines.

### RB-family knockdown is insufficient to induce *APOBEC3B* expression

RB1 is the most widely studied target of the LXCXE motif of viral proteins such as HPV E7, adenovirus E1A, and polyomavirus LTAg (29, 30, 35, 36). Due to the structural conservation of the LXCXE-binding clefts in other pocket proteins, RBL1 (p107) and RBL2 (p130) are also efficiently targeted for degradation by LTAg and truncT and may be important for A3B regulation (37–40).

The aforementioned viral proteins bind to the hypophosphorylated forms of RB1, RBL1, and RBL2, which inhibits phosphorylation by cyclin-dependent kinases (CDKs) and leads to an accelerated cell cycle in part by deregulation of E2F transcriptional activities. To investigate the roles of RB1, RBL1, and RBL2 in *A3B* regulation, a series of knockdown experiments was done with siRNAs targeting each of these factors in RPTECs and MCF10A (**Fig. 4A-B**). RT-qPCR showed that >75% knockdown was achieved for each targeted gene (upper panels in **Fig. 4A-B**). As controls, *CCNE2* was upregulated upon *RB1* knockdown and *UNG2* was upregulated by *RBL2* knockdown (lower panels in **Fig. 4A-B**). However, surprisingly, no combination of siRNAs resulted in significant *A3B* upregulation (lower panels in **Fig. 4A-B**). These results indicate that depletion of RB family members is alone insufficient to upregulate *A3B*, at least in these two different normal-like cell types where *A3B* can be induced. These results also suggest that truncT in aforementioned experiments may have at least one additional activity mediated the LXCXE motif that contributes to *A3B* upregulation.

**Figure 4.**
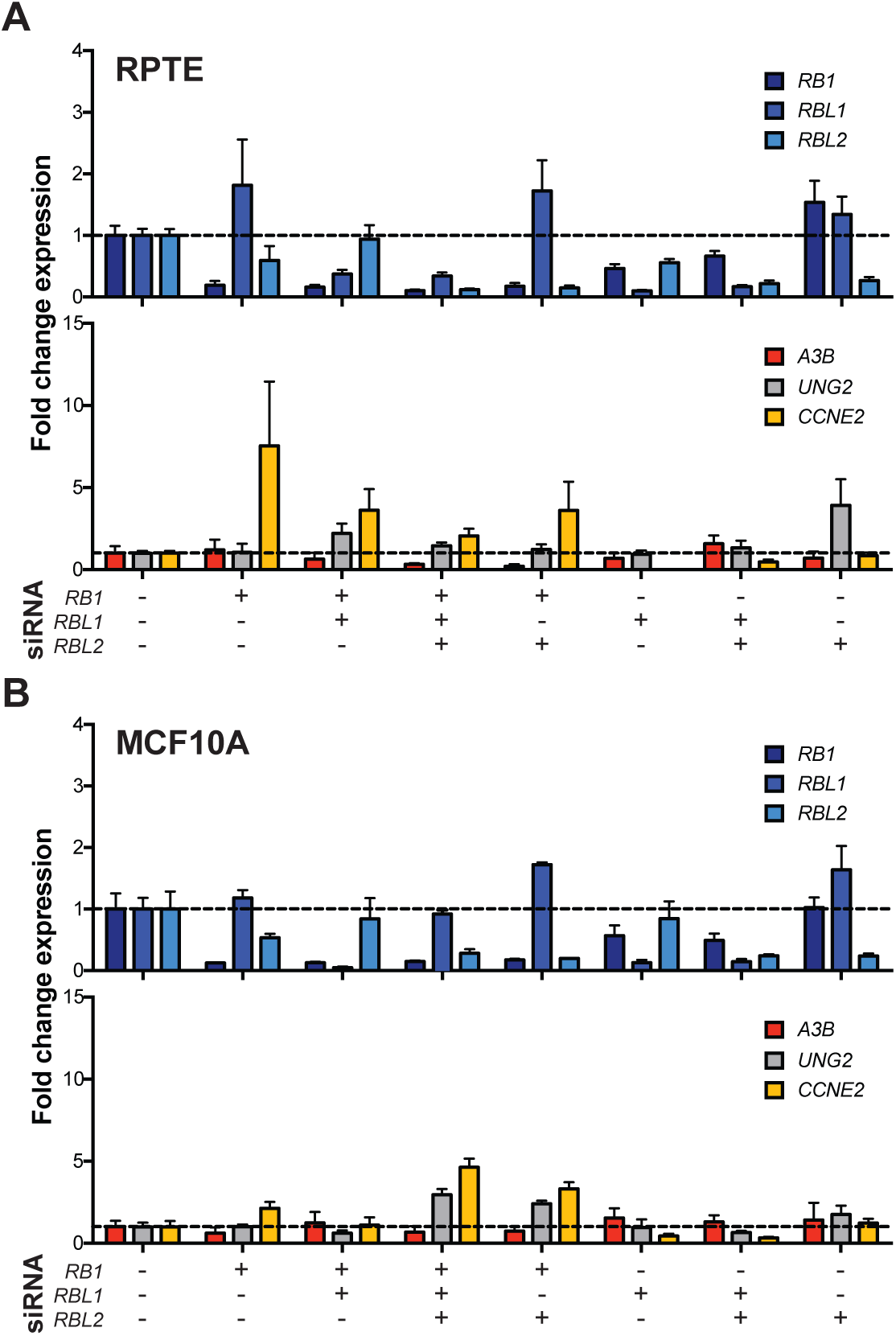
Modulation of RB family genes and A3B regulation. (A-B) RT-qPCR quantification of *RB*-family genes (top) and predicted responsive genes, *A3B*, *UNG2*, and *CCNE2* (bottom), in RPTEC and MCF10A with KD of RB-family genes.

### Pharmacological inhibition of CKD4/6 does not alter *APOBEC3B* expression

Palbociclib is a selective inhibitor of CDK4 and CDK6, which are kinases that function normally to phosphorylate RB, prevent binding to E2F transcription factors, and stimulate the expression of many genes involved in cell cycle progression (41–43). To corroborate the knockdown experiments above we treated a panel of transformed cell lines with palbociclib and quantified mRNA expression levels over time. This diverse cell panel was constructed based on *A3B* expression, ranging from low to high (44, 45), *TP53* status, and ability to phosphorylate RB. As a positive control for palbociclib efficacy we analyzed expression of *CCNE2*, which encodes Cyclin E2, promotes entry into S phase, and is a known CDK4/6-RB regulated gene (46, 47). The majority of cell lines showed a dose- and time-responsive decrease in *CCNE2* expression (**Fig. 5A**). This effect was minimal in HCC1937 and HCC1599 cells, which are known to display decreased RB phosphorylation (48, 49). In contrast, none of the palbociclib treated cell lines showed a reproducible or significant change in *A3B* mRNA expression. In addition, MCF10A cells were treated with PMA to induce *A3B* mRNA expression by the PKC/ncNF-/B pathway and, again, palbociclib had little effect (**Fig. 5B-C**, palbociclib added post- or pre-PMA addition, respectively). These results combine to indicate that the kinase activity of CDK4 and CDK6 is dispensable for *A3B* expression in multiple different cell lines.

**Figure 5.**
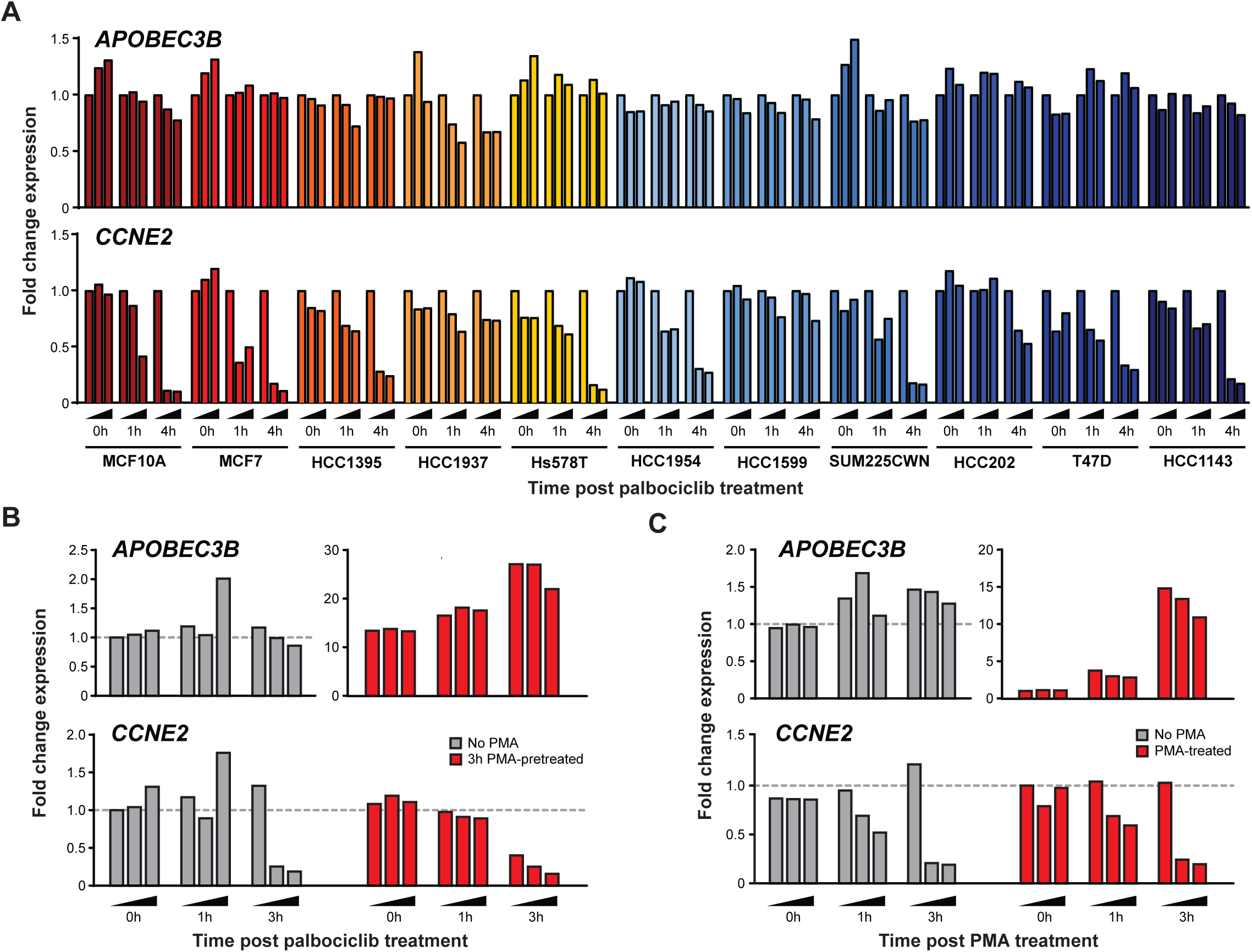
Palbociclib treatment of cancer cell lines and A3B expression. (A) RT-qPCR quantification of *A3B* and *CCNE2* expression in the indicated cell lines treated for 0, 1, and 4 hrs with increasing concentrations of palbociclib. (B) RT-qPCR quantification of *A3B* and *CCNE2* expression in MCF10A pretreated with 25 ng/ml PMA to induce *A3B* prior to treatment with increasing concentrations of palbociclib at 0, 1, and 3 hours post palbociclib treatment. (C) RT-qPCR quantification of *A3B* and *CCNE2* expression in MCF10A pre-treated with increasing concentrations of palbociclib then treated with PMA 3 hours later to induce *A3B* and evaluated at 0, 1, and 3h post PMA treatment.

### Cancer transcriptome analyses support involvement of RB-pathway in *APOBEC3B* regulation

We next used bioinformatics approaches to mine TCGA data and assess global correlates with *A3B* expression in human tumors. First, we conducted pathway analysis using all genes with significant positive correlations between *A3B* expression in TCGA breast cancer cohort. This analysis revealed that half of the top 20 significantly enriched upstream transcription factors contributing to this expression pattern are part of the CDK4/6-Cyclin D-RB-E2F axis (green-labeled genes in **Fig. 6A**). These regulators were either significantly activated or inhibited, generally corresponding with known functions, with the net outcome being accelerated cell cycling (respectively, red and blue colored bars in **Fig. 6A**). Upon closer pair-wise examination of effectors in this signal transduction pathway, *A3B* mRNA expression has the strongest positive correlations with expression of *RBL1*, *E2F1*, *E2F2*, *E2F7*, and *E2F8* (**Fig. 6B**). Lastly, we expanded this expression correlation analysis to 22 cancer types in TCGA and all 11 *APOBEC* family members. This global approach further highlighted strong correlations between *A3B* and expression of *E2F1*, *E2F2*, *E2F7*, and *E2F8*, and indicated that the association between *A3B*, this signal transduction pathway, and the cell cycle is evident in many cancer types (**Fig. 6C**). Heatmap intensities also indicated that *A3B* is the only *APOBEC* family member that correlates globally with activation of the RB-E2F axis.

**Figure 6.**
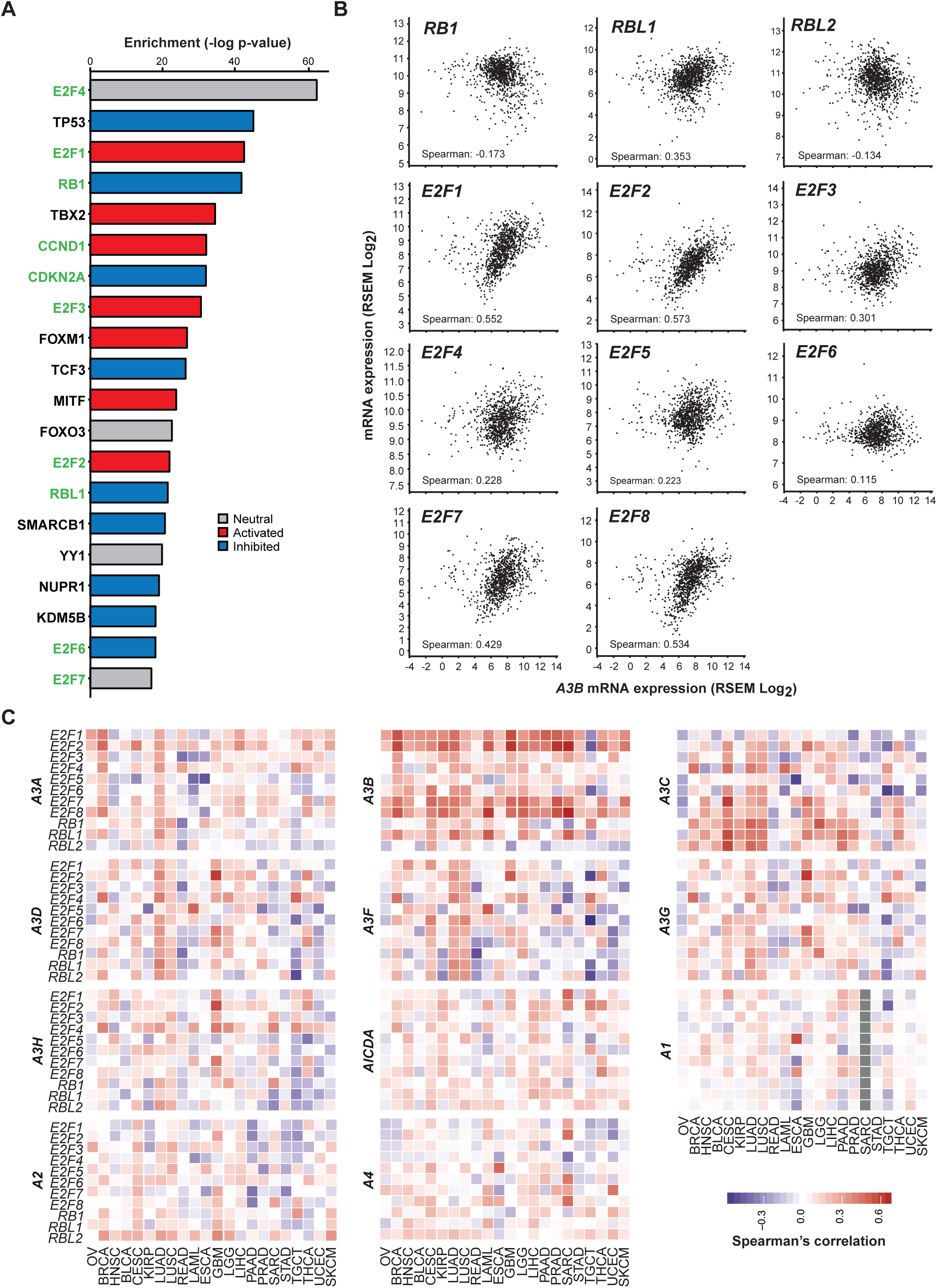
Evidence for *A3B* regulation by the RB/E2F pathway in tumors. (A) Top 20 hits from enrichment analysis of upstream transcriptional regulators of *A3B* in TCGA breast cancer with RB-pathway related genes highlighted in green. (B) Scatter plots showing the correlation between *A3B* and transcription factors in the RB pathway. (C) Spearman’s correlation coefficient values for all *APOBEC*-family members against RB pathway transcription factors across 22 cancers ordered by hierarchal clustering.

## Discussion

In this study, we investigate the mechanism of *A3B* upregulation by polyomavirus T antigen through separation-of-function mutants, genetic knockdowns, and bioinformatics approaches. First, we use high-resolution fluorescent microscopy to show that polyomavirus infection causes *A3B* upregulation and protein accumulation in the nuclear compartment. Second, we show that the LXCXE motif of LTAg and truncT, which is well known to inhibit the tumor suppressor RB, is essential for *A3B* upregulation, whereas the TP53-binding domain is dispensable. Third, genetic and pharmacologic treatments indicate that RB family members (RB, RBL1, and RBL2) and activity from kinases responsible for their phosphorylation (CDK4 and CDK6) are dispensable for *A3B* mRNA upregulation. However, bioinformatics analyses of tumor expression data show strong global correlations between *A3B* mRNA expression and expression of other genes regulated by the RB-E2F signaling pathway including several members of the E2F family of transcription factors. Taken together, these results indicate that this pathway, which is commonly modified in cancer, contributes to *A3B* gene expression and, once upregulated/activated, then *A3B* expression appears capable of continuing without additional signaling from this pathway (model in **Fig. 7**).

**Figure 7.**
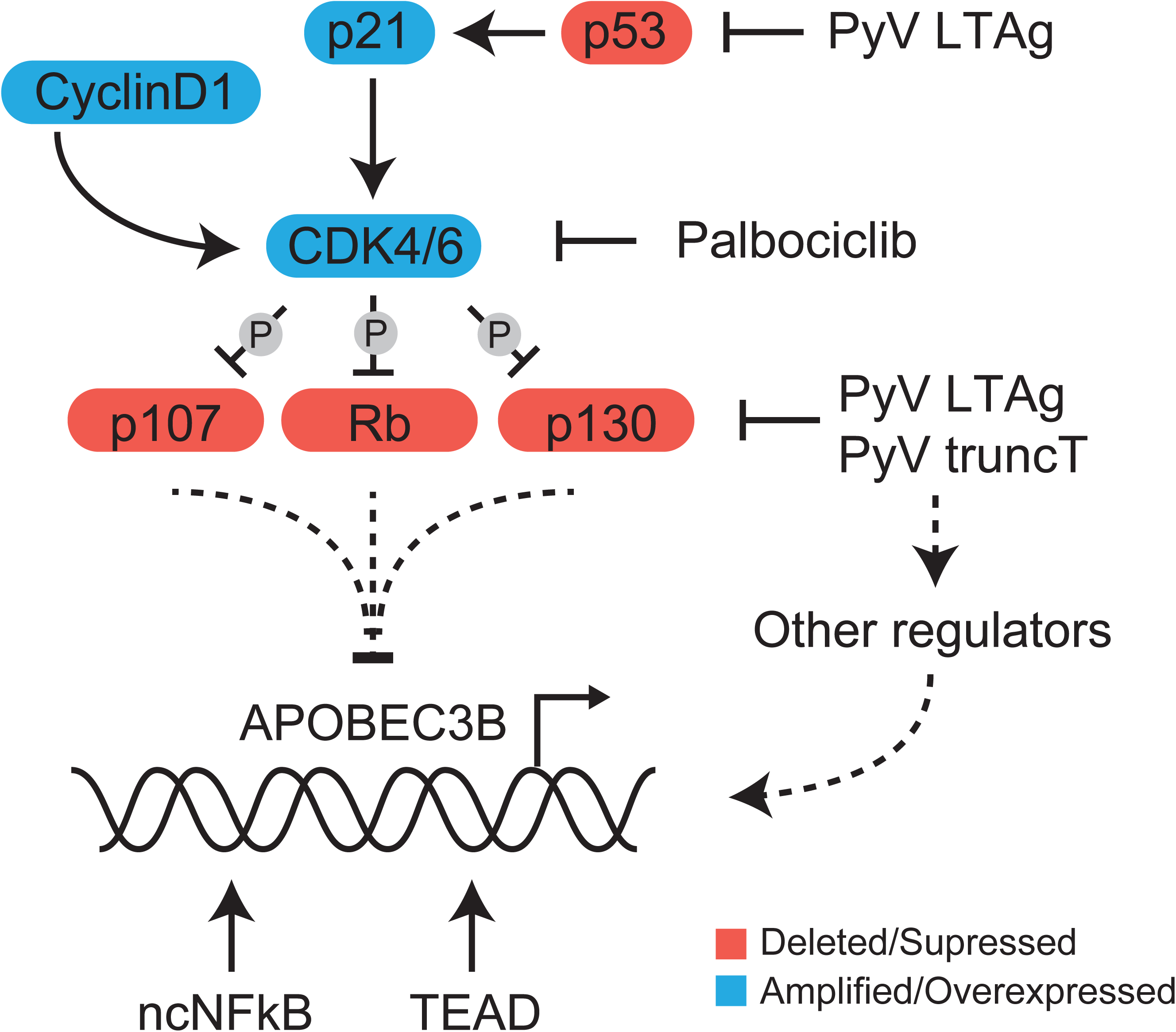
Model for *A3B* transcriptional regulation. Schematic of the cell cycle-related proteins affected by T-antigen and drug treatments used in this study with implications for *A3B* regulation. Dotted lines represent currently undefined regulatory interactions/pathways.

Our original studies with HPV and *A3B* led us to propose a model in which TP53 represses *A3B* transcription and that TP53 inactivation by the viral oncoprotein E6 or by genetic mutation results in derepression of *A3B* transcription (50). This transcription repression model is consistent with strong correlations in tumors and cell lines between *A3B* overexpression and *TP53* inactivation (44). A recent study confirmed these correlations and used pharmacologic and genetic approaches to provide further support for such a transcriptional model (33). However, three different results here do not support this TP53-mediated model for *A3B* repression. Specifically, the TP53 binding domain of BKPyV is dispensable for *A3B* upregulation, nutlin treatment has no effect on basal or induced *A3B* expression, and *TP53* knockout fails to induce *A3B* expression (**Fig. 2** and **Fig. 3**). Moreover, another recent study showed that TP53 inactivation renders cells more permissive for A3B overexpression and mutagenesis (45). Therefore, we now favor a “tolerance model” in which TP53 inactivation (genetic, epigenetic, or viral) is required for cells to be able to tolerate *A3B* overexpression. This model also explains why TP53 mutation is the only clear global correlate with *A3B* overexpression in cancer.

The RB-E2F signaling pathway is one of the most frequently mutated in cancer, which contributes to several hallmarks of cancer by deregulating the cell cycle (51). RB-inactivation is also a common target for viral genes in order to promote the survival of infected cells. The integration and continued expression of these genes in the host genome are common characteristics of virus-associated tumors. For example, HPV-associated tumors frequently have genetically conserved copies of the E7 oncogene integrated into the tumor genome that are critical for tumor development (52). These tumors also tend to have a high burden of APOBEC-associated mutations predominantly on the lagging strand, which is synthesized during the S-phase (12, 15, 53). Although some of these effects have been explained by perturbations to the TP53 pathway, other pathways have also been shown to alter *A3B* expression (23, 27, 33, 50). It is therefore not surprising in hindsight that expression of the antiviral enzyme, A3B, is also induced when the RB-E2F pathway is disrupted. Furthermore, previous studies also observed that elevated A3B expression is significantly correlated with proliferative features in breast cancer (54). Together this suggests a complex network of A3B regulation, to prevent potentially oncogenic mutations of the host genome during normal cellular replication, while activating the enzyme during abnormal cell cycling due to viral infection or cancer.

## Materials and Methods

### Cell lines, culture conditions, and lentivirus production

Primary renal proximal tubule epithelial cells (RPTEC, Lonza) were grown in REGM (Lonza), MCF10A cells were grown in MEGM (Lonza) containing penicillin (100 U/ml) and streptomycin (100 μg/ml). HuK(i)G10 cells were grown in RenaLife Epithelial Medium (Lifeline Cell Technologies) with 5% FBS, primary human choroid plexus cells were grown in EpiCM (ScienCell) on plates coated with L-glutamine and MCF7 and derivative cell lines were grown in Richter’s modification medium containing 5% fetal bovine serum, penicillin (100 U/ml), streptomycin (100 μg/ml), and 11.25 nM recombinant human insulin. All lines were grown at 37°C in a CO_2_ incubator. Lentiviruses expressing TAg, and mutants were produced in 293T cells and transduced into RPTE cells as described previously (21). BKPyV infections were conducted in MCF10A cells as described previously in RPTECs (55).

### Antibodies

The pAb416 antibody was used against BK T-antigen, which detects both Large T and truncT (28). PAB2000 was used against JC large T-antigens and mutants (56). Harris lab manufactured rabbit monoclonal antibody 10.87.13 was used to detect APOBEC3B (57). UNG2 was detected using the Abcam antibody #23926.

### RNA isolation, RT-qPCR and Western Blot

Total RNA was harvested by removal of medium and resuspension in TRIzol (Thermo Fisher), and purification was done per the manufacturer’s protocol. Reverse transcription-quantitative PCR (RT-qPCR) was used to quantify A3 transcripts as described previously (21). Protein lysates were harvested at 7 days post infection (dpi) or 3 days post transduction, quantified, and immunoblotted as described previously (55).

### Immunofluorescence

HuK(i)G10 kidney cells were seeded at 6000 cells/well in a 96-well plate. 24 hours later, infection with JCPyV was performed as previously described and then cells were collected 7 days post infection. Infected cells were incubated with EdU (Click-iT Plus EdU Alexa Fluor 647 Imaging Kit, Thermo Fisher Scientific) for 15 min and incubated with CSK buffer (10 mM Hepes-KOH, pH 7.4; 300 mM sucrose; 100 mM NaCl; 3 mM KCL; 0.5% triton x-100) (58) for 2 min on ice. Cells were then fixed in 4% PFA for 10 min followed by permeabilization with 0.5% Triton x-100 for 20 min. For EdU detection, the Click-iT reagent was added for 30 min in the dark following manufacturers protocol and washed 3x with PBS. Samples were incubated with BlockAid blocking solution (Thermo Fisher) for 1 hour at RT. T-antigen, VP1 and A3B staining were performed using the aforementioned antibodies at 1:1000, 1:1000 and 1:100 dilutions in BlockAid, respectively, overnight at 4°C followed by staining with the secondary antibodies for 1h at RT. Images were acquired on the Phenix with the confocal 63x water objective.

### siRNA and expression construct transfection

siRNAs targeting RB1 (J-003296-23, Dharmacon), RBL1 (SI02629921, Qiagen), RBL2 (sc-29425, Santa Cruz), and fluorescein conjugated non-targeting control siRNA (sc-36869, Santa Cruz) were purchased and diluted to a working concentration of 20μM. A final concentration of 20nM was used for all targets in RPTECs and 40nM in MCF10A. Reverse transfection was performed using Lipofectamine RNAiMax (Thermo Fisher) as the transfection reagent as previously described (59).

### Drug treatment

Cells were treated with 5μM nutlin (Sigma) for 24h and, after 18h of treatment, 25ng/mL phorbol myristate acetate (PMA, Sigma) was added for the final 6h prior to RNA extraction. MCF10A cells were treated only with PMA or DMSO and RNA was isolated 6h after treatment. For the palbociclib experiments, MCF10A and MCF7 cells (*TP53* WT and low *A3B* expression), HCC1937 and HCC1395 (low *A3B* expression), T47D, HCC1954, and Hs578T (intermediate *A3B* expression), and HCC1599, HCC1143, SUM-225-CWN, and HCC202 (high *A3B* expression) cells were cultured in 6-well plates (Costar 3516, Corning Incorporated) until 70% confluence. Palbociclib (S1116, Selleckchem), stored as a 5mM solution in H_2_O, was added to cells at concentrations of 0μM (H_2_O control), 0.5μM, and 2.5μM. No palbociclib was added to cells of the 0h time point, which instead was transferred to ice prior to RNA isolation. RNA was also isolated at 1h and 8 h post palbociclib administration (Total RNA Purification Kit 37500, Norgen), and cDNA was synthesized with 500ng RNA (iScript, 170-8891, Bio-Rad). RT-qPCR assays for *A3B* and *CCNE2* were performed using the C1000^TM^ Thermal Cycler (Bio-Rad). For pre-treatment with PMA, cells were treated with 0 or 25ng/ml PMA for 3 hours, followed by treatment with 0μM, 0.5μM, and 2.5μM palbociclib. RNA was isolated at 0, 1, and 3 hours, and processed as described above. Pre-treatment with palbociclib included a 0, 1, or 3 hour treatment with 0 or 25ng/ml PMA, added to cells which had been treated with 0μM, 0.5μM, and 2.5μM palbociclib 30 minutes earlier. RNA was isolated at 0, 1, and 3 hours post PMA addition, and processed as described above.

### Bioinformatics

TCGA expression data were downloaded from the Broad GDAC Firehose as of January 2016. Expression correlations against APOBEC3B by all other genes in the breast cancer cohort were calculated and significant correlates were used to determine significant activation or inhibition of upstream regulators using Ingenuity Pathway Analysis (Qiagen). All other Spearman correlations and p-values were calculated and heatmaps were plotted using the R statistical environment.

## Acknowledgements

We thank Diako Ebrahimi and N. Alpay Temiz for valuable bioinformatics discussions. This work was supported in part by Biogen and by grants from the National Institutes of Health (NIAID R01 AI123162 to MJ, NCI R21 CA206309 to RSH, and NIAID R37 to RSH). JWMM, PS, and RSH received funding from the Dutch cancer Society (Grant no. EMCR-2016-10270). Salary support for T.A.S. was provided by the Susan G. Komen Foundation, G.J.S. and J.M.M. by a Graduate Research Fellowship from the National Science Foundation, and M.C.J. by a training grant from the NCI (T32 CA009138). RSH is the Margaret Harvey Schering Land Grant Chair for Cancer Research, a Distinguished McKnight University Professor, and an Investigator of the Howard Hughes Medical Institute.

